# Visual Motion Coherence Responses in Human Visual Cortex

**DOI:** 10.1101/2021.10.18.464860

**Authors:** Andriani Rina, Amalia Papanikolaou, Zong Xiaopeng, Dorina T. Papageorgiou, Georgios A. Keliris, Stelios M. Smirnakis

## Abstract

Random dot kinematograms (RDKs) have recently been used to train subjects with cortical scotomas to perform direction of motion discrimination, partially restoring visual motion perception. To study recovery of visual perception, it is important to understand how visual areas in normal subjects and subjects with cortical scotomas respond to RDK stimuli. Studies in normal subjects have shown that Blood Oxygen Level Dependent (BOLD) responses in human area hV5/MT+ increase monotonically with coherence, in general agreement with electrophysiology studies in primates. However, RDK responses in prior studies were obtained while the subject was performing fixation, not a motion discrimination condition. Furthermore, BOLD responses were gauged against a baseline condition of uniform illumination or static dots, potentially decreasing the specificity of responses for the spatial integration of local motion signals (motion coherence). Here we revisit this question starting from a baseline RDK condition of no coherence, thereby isolating the component of BOLD response due specifically to the spatial integration of local motion signals. In agreement with prior studies, we found that responses in area hV5/MT+ of healthy subjects were monotonically increasing when subjects fixated without performing a motion discrimination task. In contrast, when subjects were performing an RDK direction of motion discrimination task, responses in area hV5/MT+ remained flat, changing minimally, if at all, as a function of motion coherence. A similar pattern of responses was seen in area hV5/MT+ of subjects with dense cortical scotomas performing direction of motion discrimination for RDKs presented inside the scotoma. Passive RDK presentation within the scotoma elicited no significant hV5/MT+ responses. These observations shed further light on how visual cortex responses behave as a function of motion coherence, helping to prepare the ground for future studies using these methods to study visual system recovery after injury.

## 2 Introduction

Visual motion perception enables us to navigate the environment and avoid collisions with obstacles. In humans, a cardinal area for motion perception is located in the posterior bank of the superior temporal sulcus in the dorsal middle temporal cortex (area hV5/MT+) (Huk, Dougherty, and Heeger 2002; Becker, Erb, and Haarmeier 2008; Helfrich, Becker, and Haarmeier 2013).

Random Dot Kinematograms (RDKs) have been used extensively to study the spatial motion integration properties of visual areas of macaques (W. Newsome and Pare 1988; Rees, Friston, and Koch 2000; Britten 1998) and humans (Rees, Friston, and Koch 2000; Muckli et al. 2002; Vaina et al. 2001; Saionz et al. 2020). RDK stimuli force the visual system to extract the global coherent direction of motion from local motion signals that have to be integrated over space and time (O. Braddick 1974; W. Newsome and Pare 1988; Scase, Braddick, and Raymond 1996; Watamaniuk, Grzywacz, and Yuille 1993) before motion direction can be perceived. The strength of the motion signal of an RDK is modulated either by changing the fraction of dots that move in the same direction (coherence) among a background of randomly moving dots (W. Newsome and Pare 1988) or by narrowing the range of directions of motion that each dot can take from frame to frame (K. R. Huxlin et al. 2009; Krystel R. Huxlin and Pasternak 2004). In translational studies, RDKs are being used for rehabilitation of visual motion perception following visual system lesions, and in particular to study the recovery of direction of motion perception following primary visual cortex (V1+) lesions (K. R. Huxlin et al. 2009, 2005; Barbot et al. 2020; Cavanaugh et al. 2019; Saionz et al. 2020). It is important to study the response of visual areas to RDK stimuli in healthy humans as well as in subjects with cortical scotomas at baseline, prior to training, in order to better understand how global motion integration is processed in patients versus normal controls. Furthermore, it is important to study RDK processing under two different conditions, i.e. when subjects perform a task related to motion discrimination versus a motion-unrelated task at fixation, as task performance is known to modulate cortical area responsiveness [[refs]].

Electrophysiological studies in macaques (Britten and Newsome 1998; Dubner and Zeki 1971; Allman and Kaas 1971; Priebe, Lisberger, and Movshon 2006; Rodman, Gross, and Albright 1989) demonstrated that the responses of middle temporal (MT) and middle superior temporal (MST) neurons are tuned to the strength of coherent motion (Heuer and Britten 2007). Specifically, the firing rate of directionally selective neurons in area V5/MT increases linearly with RDK coherence (Britten et al. 1993) and is correlated with the strength of the monkey’s direction of motion perception (W. T. Newsome, Britten, and Movshon 1989; Parker and Newsome 1998). Neurons in other visual areas show variable responses to the coherent motion of RDKs, with early visual areas including V1 tending to show suppression when RDKs are presented over a large field of view. Functional magnetic resonance imaging (fMRI) has been used to measure the responses of human visual areas to motion coherence with less consistent results. Studies in normal subjects show, as expected, higher BOLD responses for RDK stimuli with high motion coherence in area hV5/MT+, especially when these are presented over a large field of view (Rees, Friston, and Koch 2000; Oliver J. Braddick et al. 2001; Becker, Erb, and Haarmeier 2008). However, other reports are conflicting, reporting hV5/MT+ BOLD responses do not depend on (Smith et al. 2006; Beer et al. 2002) or even decrease with motion coherence (McKeefry et al. 1997; Previc et al. 2000). The response of human visual areas other than hV5/MT+ as a function of coherence is less clear. Rees et al. (Rees, Friston, and Koch 2000) report the BOLD response is linear as a function of RDK motion coherence in area hV5/MT+, while higher areas (‘kinetic occipital’ (KO), V3A, frontal cingulate gyrus) tend to show either non-linear U-shaped responses or negative correlation.

Human fMRI studies typically measure the responses of large populations of neurons with diverse properties (Tootell et al. 1995; Zeki et al. 1991) rather than single cells (Kwong et al. 1992) tuned to the direction of motion, and may therefore be more sensitive to the overall state of adaptation and to the brain state of the subject while they are performing a task. Task performance, in part through attention, has been known to modulate neural activity in the visual cortex (Azzopardi and Cowey 2001), specifically in the dorsal stream including hV5/MT+ (Tootell et al. 1995; Schoenfeld et al. 2007; Saenz, Buracas, and Boynton 2002; Heeger et al. 1999), and it is natural to ask whether the linear response of area hV5/MT+ to motion coherence is also affected. To explore whether RDK BOLD responses depend *1)* on the state of adaptation, i.e. the baseline visual stimulus condition from which RDK stimuli of various global motion coherence strength are presented (Tolias et al. 2001), and *2)* on direction of motion discrimination task performance, we made two protocol modifications: ***1)*** We measured RDK responses as a function of motion coherence when the subject was performing a direction of motion discrimination task versus when they were simply fixating. ***2)*** We measured the response to coherent motion starting from a baseline state elicited by an RDK of zero coherence rather than a uniform gray screen. Starting from this 0% motion coherence baseline allowed us to map responses selective to the spatial integration of coherent motion signals, as opposed to responses induced by the combined changes in local luminance and motion contrast.

Our results under simple fixation starting from a 0% coherence baseline condition corroborated the result that area hV5/MT+ activity increases as a function of coherence in agreement with (Rees, Friston, and Koch 2000). However, when subjects performed the RDK direction-of-motion discrimination task coherence dependence was essentially abolished. A similar result was obtained in area hV5/MT+ of subjects with dense cortical scotomas performing direction of motion discrimination for RDKs presented inside the scotoma, whereas passive RDK presentation within the scotoma elicited no significant hV5/MT+ responses. Our results complement the existing literature and help to inform the design of future RDK mapping experiments to study how visual areas reorganize following visual motion perception rehabilitation in subjects with cortical scotomas.

## 3 Material and Methods

### 3.1 Human Subjects

The human experiments consist of three separate RDKs studies.

#### Study A

Six healthy subjects with no history of psychiatric or neurological problems were recruited. Two subjects were excluded from the analysis because of significant head motion in the MR environment (>5 mm), which could not be adequately corrected offline.

#### Study B

For the second study, six (6) more healthy participants who fulfilled our inclusion criteria underwent the experimental procedure.

All the participants for both studies were recruited at the Core for Advanced MR Imaging at f Baylor College of Medicine (BCM)

#### Study C (patients and controls)

*Patients:* seven subjects (27-64 years old, 3 females – 4 males) with visual cortical lesions participated in our study. Six of them were recruited at the Max Planck Institute for Biological Cybernetics in Tübingen (MPI) and one at the Core for Advanced MR Imaging of the Baylor College of Medicine (BCM). *Controls:* Six healthy subjects were recruited as control subjects. Four of them were scanned at MPI and two at BCM.

All subjects had normal or corrected-to-normal visual acuity. Experiments were done with the approval of the IRB committees of Baylor College of Medicine and the Regierungspräsidium of the Max Planck Institute for Biological Cybernetics, Tübingen, Germany.

##### 3.1.1 MRI Scans

###### For studies A and B

Functional and structural scans were performed at the Core for Advanced MR Imaging at BCM, using two 3.0 Tesla MRI scanners (Siemens Ltd, Erlangen, Germany): Allegra and TIM Trio, both equipped with a quadrature 12-channel coil. T1-weighted high resolution (MPRAGE) scans, for about 7 min each, were acquired twice for SNR reduction (TR=1900ms, TE=2.26ms, matrix size = 256×256×192, flip angle=9°, spatial resolution Trio/Allegra=0.5×0.5×1.0mm^3^/0.96*0.96*1.0mm^3^). Blood-oxygen-level-dependent (BOLD) images were registered onto the anatomical images, which were used to (i) co-register the functional scans to the anatomy of each subject for each session, and to (ii) segment each subject’s anatomical data into white and gray matter. Whole brain T2*-weighted BOLD images were acquiring using the single-shot echo planar imaging pulse sequences were acquired covering the entire brain (repetition time [TR] = 2000ms, echo time [TE] = 40ms, matrix size = 64 × 64, voxel size = 3.28×3.28×3.28 mm^3^, flip angle = 90 degrees, number of slices Trio/Allegra=28/29).

#### For study C

The scans were performed on a 3.0 Tesla Siemens Prisma (at MPI) and Trio (at BCM) (Siemens Ltd, Erlangen, Germany). For each participant (patient or control subject), two T1-weighted anatomical images were acquired (*<u>Prisma</u>*: voxel size=1×1×1 mm^3^, matrix size= 256×256×192, flip angle= 9°, TR= 2300ms, TE =2.98ms, TI=1100ms; *<u>Trio</u>*: voxel size= 0.5×0.5×1.0mm^3^, matrix size= 256 ×256, 192 partitions, flip angle= 9°, TR= 2600ms, TE =3.5ms, TI=1100ms). BOLD images were acquired using gradient echo planar sequences with 29 (MPI) and 30 (BCM) contiguous 2.6 and 2.5mm-thick slices respectively, covering the whole brain (TR= 2000ms, flip angle=90°, matrix size = 64 × 64, Prisma/Trio TE= 35/30ms, Prisma/Trio voxel size = 3×3×2.6/3×3×2.5mm^3^). For each subject five functional scans were acquired, each consisting of 131 image volumes.

##### 3.1.2 Stimulus presentation

###### Studies A and B

Stimuli were projected under photopic conditions onto a rear-projection translucent acrylic screen (DaTex, Da-Lite Corp.) via a NEC GT2150 projector (2500 ANSI Lumens, 1600 × 1200 resolution, 120Hz) controlled by a Macintosh computer, and seen through an oblique mirror mounted on the MR head coil. The active visual field subtended ∼15^0^ in radius. When necessary, subjects were corrected for optimal accommodation using magnet-compatible glasses.

###### Study C

At the MPI, for stimulus projection we used MRI compatible digital goggles (VisuaStim, Resonance Technology Company, Inc, Northridge, CA, USA), with FOV=30° (horizontal) and 22.5° (vertical), resolution= 800×600, mean-luminance 5.95 cd/m^2^. An infrared eye tracker was used to record eye movements (iView XTM, SensoMotoric Instruments GmbH). At the BCM, the stimuli were projected on the same screen as described under studies A, B,.

All stimuli were presented under photopic conditions, the dots being dark on bright background to minimize scattering and were generated using MATLAB (MathWorks), and the psychophysics toolboxes, Psychtoolbox [(Kleiner et al. 2007) http://psychtoolbox.org] and Vistadisp, an open toolbox (VISTASOFT, Stanford, https://github.com/vistalab/vistadisp).

#### 3.1.3 Stimulus Paradigms

##### Retinotopic Mapping

###### Studies A&B

Retinotopic visual field maps were obtained using phase-encoded retinotopic mapping (Engel, Rumelhart, and Wandell 1994) with 45^0^ wedges and 1.5^0^ concentric rings, and the borders of the early visual field areas determined according to (Wandell, Dumoulin, and Brewer 2007). Stimuli were circular with a maximum radius of 12^0^. A full wedge cycle was completed in 36s, with a total of 6 cycles per scan (216s). For rings, the pattern of the stimulus was moving in repeating cycles from the center to the periphery through 8 expansions of 32s each (256s). Both rings and wedges had flickering checkerboard patterns at 2Hz, spatial frequency ∼1c/deg and 100% contrast and were acquired with TR 2s. Two to five scans were performed in each stimulus condition (wedge and ring), depending on the subject.

###### Study C

The stimulus consisted of moving square-checkerboard bars (100% contrast) within a circular aperture with a radius of 11.25° around the fixation point. The bar width was 1.875° and travelled sequentially in 8 different directions, moving by a step half of its size (0.9375°) every image volume acquisition (TR=2 seconds). The subjects’ task was to fixate on a small dot in the center of the screen (radius: 0.0375°; 2 pixels) and respond to the color change (red to green) by pressing a button. The color was changing randomly with a frequency of one every 6.25 seconds.

##### Functional Localizer for hV5/MT+

Area hV5/MT+ was identified using 18s blocks that alternated between moving (100% coherence) and stationary dot patterns. To reduce adaptation, the direction of the moving dots changed by 45 degrees counterclockwise. Each scan consisted of either 6 or 8 blocks of moving and static dots, depending on the subject. Each scan’s moving and stationary dot patterns were repeated six to eight times during each functional localizer scan. A total of 48 blocks of moving and static dots were acquired for each subject.

Please note that normal retinotopic mapping based on clusters of activity at the expected anatomical locations was also sufficient for identifying hV5/MT+ and provided a second method for ensuring accurate hV5/MT+ identification. No localizer was used for the scans in Study C.

##### Motion Coherence Paradigm

**Study A** (subjects did not perform direction of motion discrimination)

###### Task

During study A, the subjects were instructed to perform a fixation task that did not involve direction of motion discrimination while passively viewing full-field RDKs with different motion coherence. A small dot of 0.15-degree radius was displayed at the center of the visual field to serve as a fixation mark. The color of the fixation point changed between green and red at random times and the subjects were required to report the color change by button press. All subjects responded correctly for > 96% of fixation color changes, with average response time < 0.6 s.

###### Stimulus

Dynamic random dot kinematograms (RDKs) were generated within a circular aperture (12-degree radius) using the method described in Newsome and Pare (W. Newsome and Pare 1988). RDK dots were dark, at a density of 2 dots/degree^2^ and radius of 0.1 deg and were presented on a gray isoluminant photopic background to minimize light scattering. The random dot pattern was refreshed every 50ms. A random fraction of the dots was displaced by 0.2° in the same direction (left or right) at a rate of 4°/second, while the remaining dots were replaced by the same number of new dots at random positions. The percent of dots moving in the same direction, i.e the strength of coherent motion signal, varied between 12.5%, 25%, 50%, 100% in a randomly counterbalanced fashion. Each coherence was presented for 10s interleaved with 30 sec of baseline stimulus at 0% percent coherence (no correlation amongst dots). All coherence levels appear for the same total time within each scan. To help subjects maintain a stable level of adaptation to the baseline (0% coherence) stimulus, the 0% coherence stimulus remained on during the whole RDK experiment, except when stimuli with non-zero coherence levels were displayed. Each scan either consisted of 2 blocks of each of 4 pseudo-randomly interleaved coherence levels (12.5%, 25%, 50%, 100%; subjects 1-3), or 4 blocks of each of 3 pseudo-randomly interleaved coherence levels (25%, 50%, 100%; subject 4). Eight (for subject 4) or ten (for subjects 1-3) scans were performed in a single session (fig. 1A)

**Figure 1:**
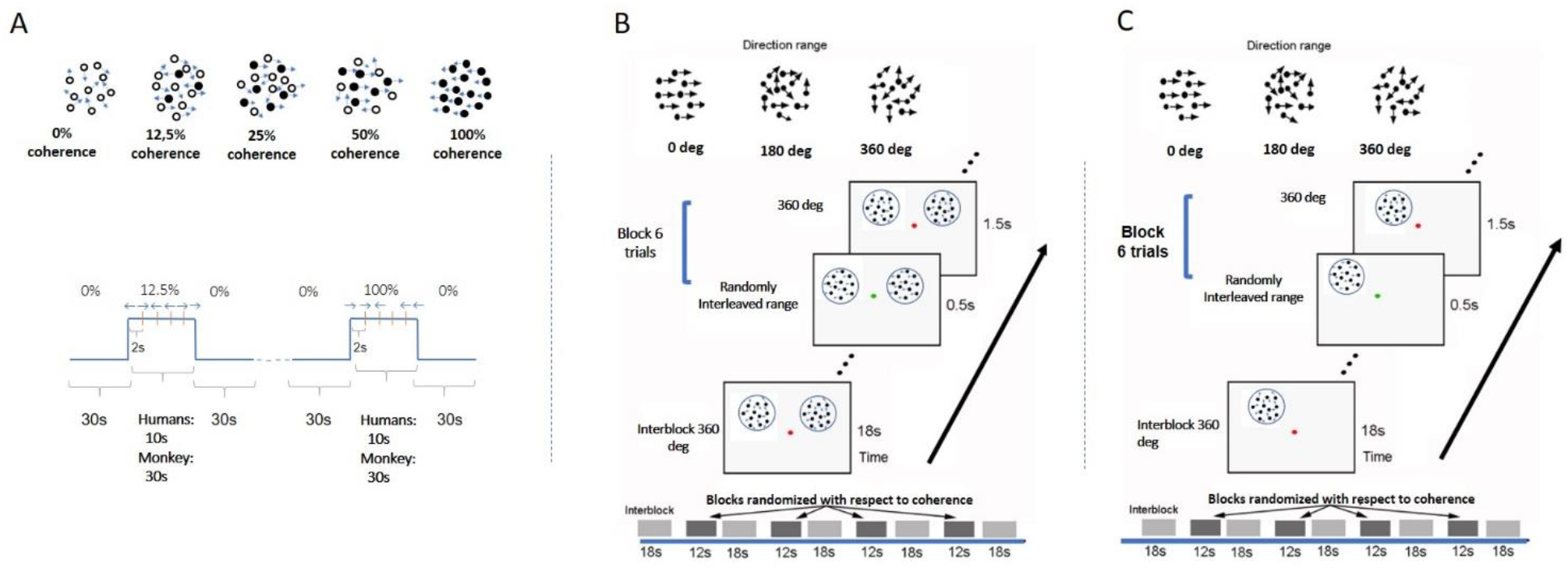
Experimental design. ***A:*** Paradigm used in study A. Each motion coherence was presented for 10s interleaved with 30 seconds of 0% coherence (no correlation amongst dots). All coherence levels appear for the same total time in each scan. Each scan either consisted of 2 blocks of each of 4 pseudo-randomly interleaved coherence levels (12.5%,25%,50%, 100%) or 4 trials of each of 3 pseudo-randomly interleaved coherence levels (25%, 50%, 100%). Within each block with non-zero coherence, the global direction changed (left vs right) every 2s to minimize adaptation. A similar paradigm was used for the monkey experiment except RDK duration was 30s for all coherences. Black dots represent the group of dots moving in the same direction. ***B:*** RDKs for study B were presented at bilaterally symmetric locations (see methods). Coherence alternated between 18sec of baseline of 360^0^ direction-range (no global coherence) followed by 12 sec of RDK coherence selected from levels (0^0^, 180^0^, 288^0^, 360^0^). All coherence levels were pseudo-randomly interleaved and balanced across blocks and scans. Within blocks coherent RDKs were presented for 0.5sec for 6 trials (see methods) and directions of motion changed every 2 sec, with left and right directions appropriately balanced but presented at random sequence (so subjects could not predict the direction). Subjects were fixating and cued by a change in the color of the fixation spot to report the direction of the stimulus in one hemifield. ***C*:** A similar paradigm to B except now RDKs were presented unilaterally. Motion discrimination versus no-motion-discrimination sessions were interleaved in the scanner.

**Study B** (bilateral stimulation; unilateral motion discrimination task performance)

###### Task

During study B, two RDKs were presented at symmetric positions in the left and right visual fields (fig. 1B), respectively, while the subjects fixated at a central spot and performed a motion direction discrimination task on the right RDK as instructed. Study B was carried out to investigate whether performing an RDK-related task changed the shape of the BOLD response profile as a function of motion coherence.

###### Stimulus

RDK stimuli for this experiment were derived according to (K. R. Huxlin et al. 2009; Krystel R. Huxlin and Pasternak 2004). The parameters were identical to the ones described above, except now each dot moves following the same rule and the global motion strength is modulated by means of the choice of the direction range of each dot (K. R. Huxlin et al. 2009; Cavanaugh et al. 2015; Saionz et al. 2020). In this design, each dot randomly “picks” a direction evenly distributed around a central direction (left or right moving). For example, when the range of motion for each dot is 360 degrees (2π), the net global directional motion signal is zero. On the other extreme, when the range of motion is 0 degrees, all the dots move toward the same direction and net global motion is full strength.

The stimuli were two RDKs 6-degree in diameter, one (task-relevant) centered at (5,4) and the other (task irrelevant) at (−5, 4), or vice versa. A dot of 0.2 degree in diameter was presented at the center to serve as a fixation spot as well as instruction cue. Each run consisted of alternating passive fixation period (18 s) and active motion discrimination period (12 s) in the right visual field. During the *active motion discrimination period*, the fixation spot changed its color from red to green every 2 sec, with green indicating the occurrence of global motion (leftward or rightward) that lasted for 0.5 sec. The subjects had to report the direction of the perceived motion in the RDK presented in their right visual field by button pressing. For each 12 sec period, the motion strength presented in each trial remained the same whereas the direction of motion varied randomly (leftward or rightward in balanced fashion). The direction of motion in the left and right RDKs were uncorrelated but the motion strength (coherence) in both presented RDK patches was the same. Four motion strength levels were tested (360°, 288°, 180°, 0°). During the *passive fixation period (RDKs at 0% motion coherence)* the fixation spot remained red and subjects were instructed to passively fixate only, while the stimuli varied identically to the active case. For each scan, each motion strength level was repeated twice, making the total duration of the run 258 sec. Each subject underwent 10 scans per session, resulting in 20 repeats per motion strength level.

**Study C** (unilateral stimulation; motion discrimination versus fixation task)

###### Task

Study C, consisted of an active and a passive task. For both, the subjects had to fixate a dot at the center of the screen. In the *fixation task*, the color of the fixation dot changed from red to green at random intervals and the participants had to respond any time there was a change. In the *motion discrimination task* experiment, participants were still instructed to fixate the dot but this time they had to use their peripheral vision to assess the direction of motion of the RDK stimulus. Whenever there was a change in the color of the central dot, they were asked to indicate the motion direction of the stimulus (left or right).

###### Stimulus

The RDK stimulus used here was similar to study B, except that RDK dots were moving slightly faster at 10°/sec (vs 4°/sec in B); the RDK aperture was presented unilaterally, either at the left or right upper quadrant of the visual field (counterbalanced), centered at 4 degrees from the vertical meridian and 3 or 4 degrees above the horizontal meridian with an aperture diameter of 4 or 5 degrees respectively (for the controls) (fig. 1C).

For each *patient* the aperture location and diameter were adjusted in order to fall within their visual field scotoma (S15: center = [5°, 4°], diameter = 5°, S29: center = [5°, 4°], diameter = 6°, S12: center = [5°, 4°], diameter = 7°, V1003: center = [5°, 4°], diameter = 5°, S07: center = [7°, 4°], diameter = 5°, S04: center = [8°, 4°], diameter = 3°, S02: center = [3°, 3°], diameter = 5°, where [x, y] = x°from the vertical meridian and y^0^ from the horizontal meridian). Four motion coherence levels (360°, 288°, 180°, 0°) were presented as in study B, where again direction of motion range at 360° corresponded to the baseline condition (no coherence).

For the *control subjects*, the aperture was presented either at the left or right upper quadrant of the visual field and was centered at 4°from the vertical meridian and, depending on subject, 3°or 4°from the horizontal meridian with an aperture diameter of 4°or 5°respectively.

##### Patients’ anatomical lesions and visual field tests

All patients had ischemic or hemorrhagic strokes 7 to 10 years prior to enrollment. These resulted in dense (visual sensitivity<-20dB) homonymous visual field deficits within either one or two quadrants of the visual field. MRI anatomical images confirmed the location and extent of the injuries (fig. 2A). In more detail, patient S02 had a left temporal and partial parietal optic radiation injury due to an infarct of the right mid-posterior temporoparietal lobes. Patients V1003 and S29 had right and left homonymous hemianopia respectively. Patient S04 had a lesion in the right hemisphere that involved part of the foveal V1v, V2v and V3v, resulting in a dense left upper visual field quadrant scotoma. The lesion of patient S07 was located in the left inferior calcarine cortex, involving areas left V1v and V1d, left V2v, left V3v and left V4, resulting in a right homonymous superior quadrantanopic defect. Patient S12 had a lesion in the right inferior calcarine sulcus, involving part of area V1 and visual areas V2v and V3v. Finally, patient S15 had a left temporal optic radiation infarction causing a dense right upper visual field quadrant defect. V1 gray matter remained intact, but a part of it lost its input.

**Figure 2:**
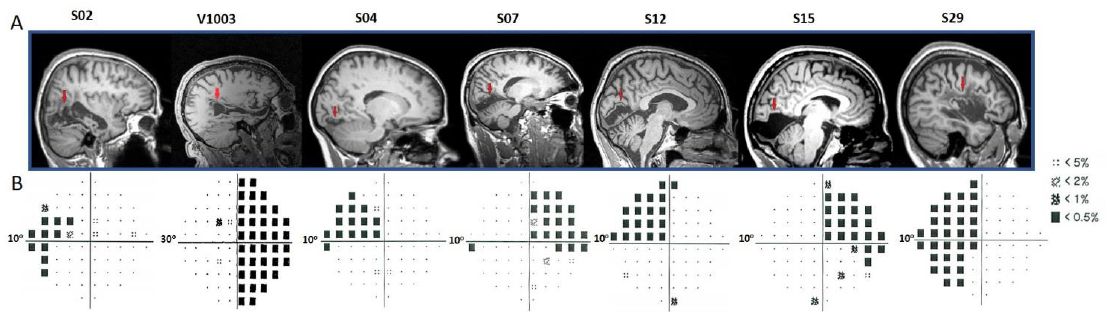
Humphrey’s Visual Field perimetry test and lesions’ location. ***A:*** Sagittal anatomical planes of patients’ brains. The red arrows point to the location of individual’s anatomical lesions. ***B:*** Subjects S02, S04,S07,S12,S15 and S29, underwent a 10-degree (10-2) and V1003 a 30-degree (30-2) Humphrey’s perimetry test. The black squares in the pattern deviation probability plots correspond to p<-20dB. Dotted squares correspond to <-10dB of visual sensitivity. The rest of the locations (small black dots) indicate the non-affected (normal) visual field.

Patients S02, S04, S07, S12, S15 and S29 visual field defects were assessed with a Humphrey type (10-2) visual field test (Beck, Bergstrom, and Lighter 1985; Trope and Britton 1987) with a (low photopic) background luminance level of 10 cd/m^2^ (fig. 2B). The visual field defects of these patients were also verified using a binocular semi-automated 90° kinetic perimetry obtained with the OCTOPUS 101-perimeter (HAAG-STREIT, Koeniz, Switzerland) (Hardiess et al. 2010). Patient V1003 underwent a Humphrey type (30-2) visual field test.

### 3.2 Monkey experiment

We mapped motion coherence responses in area V5/MT+ of one rhesus macaque that was trained to fixate. Here instead of the BOLD signal, we measure modulations in cerebral blood volume (CBV) as function of RDK motion coherence. To this end, before each scanning session we injected the monkey with 10mg/kg MION (Monocrystaline Iron Oxide Nanoparticle) (Leite et al. 2002; Leite and Mandeville 2006). Whole brain images (TR=2000ms) at 1mm isotropic resolution using a 4-channel phased array surface coil, with the AC88 gradient insert to increase spatial resolution, were acquired at 3T (Siemens Trio at Charlestown Facility, Massachusetts General Hospital, Boston MA). Fifty slices were acquired using a GE-EPI-T2* functional imaging sequence at TE/TR =19/2000ms, 96×84 matrix, 90°flip angle. The monkey was required to fixate, while its eye movements were monitored, using an ISCAN Infrared eye tracker (ISCAN, Burlington, MA).

#### Stimulus

An RDK was constructed as described above (study A) according to the method of Newsome and Pare (W. Newsome and Pare 1988) with dot density of 1.7/deg^2^ and dot size (radius) 0.1-degree. This was presented through a central field of view ring-like aperture extending from 1 to 12-degrees. The monkey maintained fixation within a 2×2 degree fixation window. Dots were dark on a bright background to minimize scattering. Coherent dot displacement was 0.2 degrees every 50ms in the horizontal direction while the direction of the dots was reversed every 5s to minimize adaptation. A 0% coherence background of RDKs was presented for 30s, alternating with 30s of RDK stimuli of different motion coherences (12.5%, 25%, 50%, 100%), pseudo-randomly interleaved in a block design (fig.1A). The mean percent BOLD signal modulation above the 0% coherence condition was computed for the retinotopically corresponding portions of the V1, V2, V3, V3A, V5/MT, FST and MST areas.

## 4 Analysis

### Studies A & B

FMRI data were preprocessed in AFNI (Cox 1996) and then analyzed with custom MATLAB codes. Segmentation of gray from white matter was performed and cortical surface reconstruction was carried out with FreeSurfer (http://surfer.nmr.mgh.harvard.edu/) using the high resolution *T*_1_ weighted images obtained in the Trio/Prisma scanners. The surface reconstruction algorithm removed extracerebral voxels via a skull stripping routine, which results in an intensity-normalized image. Raw functional data were first corrected for slice acquisition time difference and then motion correction was performed to align all EPI images to the EPI image acquired closest to the *T*_1_ weighted images. The fMRI signal time course was detrended to remove slow linear drifts of the fMRI signal. For retinotopic mapping and hV5/MT+ localization, data from scans under the same stimulus condition were averaged together after preprocessing and before data analysis in MATLAB. The averaged time-series were then mapped to the smoothed white matter surface using 3dVol2Surf in AFNI, which generates the value for each surface node by transforming original-to standard-mesh surfaces (Saad and Reynolds 2012). The reference hemodynamic response function used to convolve the stimulus profile is a Gamma function *t*^*b*^ exp(*-t/c*), with *b*=8.6 and *c*=0.547s.

*Statistical analysis* was performed on the fMRI data in the RDK experiments using a generalized linear model (GLM) approach. For each coherence level, a regressor was generated by convolving the stimulus profile of that coherence with the hemodynamic response function above. A third order polynomial was used to model slow baseline shift. The data from different scans were concatenated for the GLM analysis. The model parameters were estimated using the 3dREMLfit program in AFNI, which uses an ARMA(1,1) model to estimate the correlation structure in noise. After GLM analysis, regions of interest (ROIs) were defined as clusters with *<u>corrected</u>* p < 0.05 across all coherence levels, and that overlap with different visual areas as determined via retinotopy, was identified. We calculated BOLD time series averages in identified regions of interest (ROIs) as follows: (i) The mean ROI time series was computed by averaging the BOLD signal across the voxels belonging to each ROI and over trials presented at the same coherence level; (ii) The baseline BOLD signal was computed from the last five time points of the baseline stimulus, plus the first time point after the transition (before the BOLD signal has had time to rise)*Retinotopic Mapping and MT localizer:* For retinotopic mapping, the measure of coherence^1^ of the average time course at the stimulus frequency is used as a measure of the BOLD response strength (VISTASOFT, Stanford, https://github.com/vistalab/vistadisp). To locate activated voxels in hV5MT+ localizer scans, we calculated the correlation coefficient between the time-courses of the BOLD signal from each voxel and the stimulus time-course, and assessed significance using the t-test. Voxels with significantly different t values from zero define area hV5/MT+. Putative area MST can be identified as the voxels activated by both contra-lateral and ipsilateral RDK stimuli, but largely overlaps with area hV5/MT, so we designate both areas together as hV5/MT+.

### Study C

The functional images were corrected for motion in between and within scans (Nestares and Heeger 2000) and aligned to the high-resolution anatomical volume using a mutual information method (Maes et al. 1996). We performed the preprocessing steps, in MATLAB using the VISTASOFT toolbox (https://github.com/vistalab/vistasoft). We fitted a general linear model (GLM) to the time course of each voxel to estimate the contribution of each direction range stimulus tested to the time course. The four conditions tested (direction range: 0°, 180°, 288°, 360°) were then contrasted against the baseline (interblock, direcation range: 360°) to estimate the dependence of each voxel on coherence. Only those voxels for which the linear model explained more than 3% of the variance in the data were retained. This threshold was set after measuring the mean explained variance during the passive task in a non-visually responsive area by selecting a region of interest (i.e. a sphere of 1cm diameter) from the lower medial prefrontal cortex and setting the value of the threshold at 3 standard deviations above the mean. In a region of interest, the percentage signal change was calculated by averaging the beta weights of each predictor for each voxel. *Retinotopy:* We identified area hV5/MT+ using the population receptive field (pRF) mapping method (Dumoulin and Wandell 2008). In short, the implementation of the pRF model is a circularly symmetric Gaussian receptive field in visual space. The center and radius of the pRF are estimated by fitting the BOLD signal responses to estimated responses elicited by convolving the model with the moving bar stimuli. We retained only those voxels in these visual areas, for which the topography explained more than 12% of the variance. This threshold was set after measuring the mean explained variance (6% ± 2%) in a non-visually responsive area by selecting a region of interest (i.e. a sphere of 1cm diameter) from the lower medial prefrontal cortex and setting the value of the threshold at 3 standard deviations above the mean.

### Reconstruction of the Lesioned Hemisphere method

Analyzing the functional data of patients with cortical visual lesions can be tricky because the lack of cortical tissue in the location of the injury. To overcome this, we used a method we developed earlier and it is described in more detailed in Papanikolaou et al. (Papanikolaou et al. 2019) in order to create a ‘hybrid’ hemisphere. In brief, the method uses information from the healthy hemisphere in order to reconstruct the damaged part.

## 5 Results

### Normal Subjects

#### RDK stimulus presentation without motion discrimination (study A)

We measured the average BOLD signal modulation across four levels of motion coherence (12.5%, 25%, 50%, and 100%) in areas V1, V2, V3, V4, and hV5MT+ as a function of time following a transition from 0% coherence. RDKs were generated using the Newsome and Pare (W. Newsome and Pare 1988) method. Subjects were asked to fixate and report the change of color of a dot at fixation (see methods), performing >96% correct in this task. As expected, BOLD signal in area hV5/MT+ (fig. 3B-C) showed strong statistically significant coherence dependence (p<0.01, 1-way ANOVA over coherence across subjects, each measurement reflecting the mean BOLD response amplitude averaged across trials at the same level of coherence for each subject). Area hV5/MT+ BOLD response increased with coherence, reaching ∼0.5% above baseline at 100% motion coherence. Area **V3A** was also significantly modulated by coherence (fig. 3B; p < 0.006, 1-way ANOVA), showing a strong increase in BOLD signal intensity at 100% motion coherence, and weaker responses at lower motion coherence levels. In contrast, areas **V3** and **V4** were not significantly modulated by motion coherence levels (p=0.66 & p=0.2 respectively, 1-way ANOVA). As reported before (Oliver J. Braddick et al. 2001) the BOLD response in area **V1** decreased at higher motion coherences compared to the 0% coherence baseline. Specifically, following a transition from 0% to 100% coherence, the BOLD signal in area V1 decreased reaching minimum ∼ 10s after the transition (fig. 3B_bottom; p <0.01 1-way ANOVA). Area **V2** showed a similar trend that did not reach significance (fig. 3B; p=0.09, 1-way ANOVA). These results were qualitatively similar to results obtained in monkey visual cortex during passive fixation, using cerebral blood volume (CBV) imaging with MION (Smirnakis et al. 2007; Mandeville and Marota 1999). Monkey V5/MT showed a clear monotonic increase of CBV signal as a function of coherence (fig. 4). Area MST behaved similarly. CBV vs motion-coherence profiles computed for areas V1, V2, V3, V3A, FST were in general less sensitive to coherence, displaying similar features as the corresponding areas in the human.

**Figure 3:**
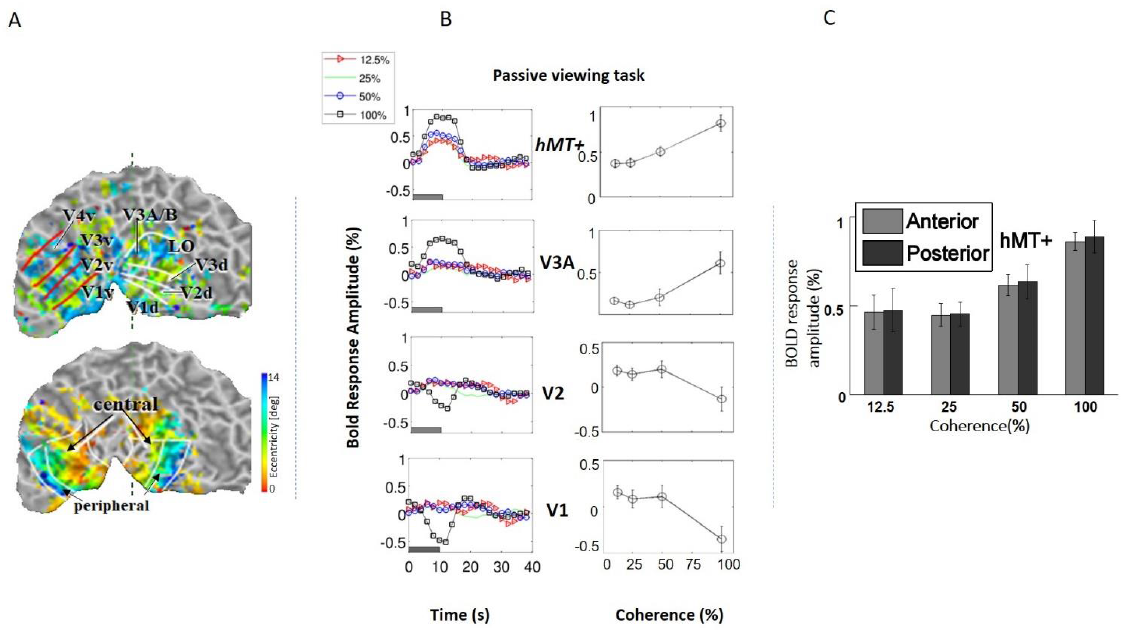
***A***. *<u>Top:</u>* example of a retinotopic map presented on the flattened visual cortex of a subject. The lines define the borders between visual areas. *<u>Bottom:</u>* example of an eccentricity map. The foveal (central) and peripheral ROIs in V1, V2 and V3 are indicated by the black arrows. ***B***. *<u>Left</u>*: Average BOLD signal intensity responses as a function of time in 4 subjects across four motion coherence levels, (12.5%, 25%, 50%, 100%) in visual areas V1, V2, V3A, hV5/MT+, while fixating (no motion discrimination task). The baseline BOLD activity elicited by 0% coherence was subtracted (see methods). *<u>Right</u>*: BOLD response amplitude as a function of motion coherence (see methods) in the passive viewing condition, across n=4 subjects. Error bars represent standard error of the mean. ***C***. Selecting the anterior and posterior voxels of hV5/MT+ did not show any difference in coherence dependence. Therefore, we grouped them together for analysis as the hV5/MT+ complex.

**Figure 4:**
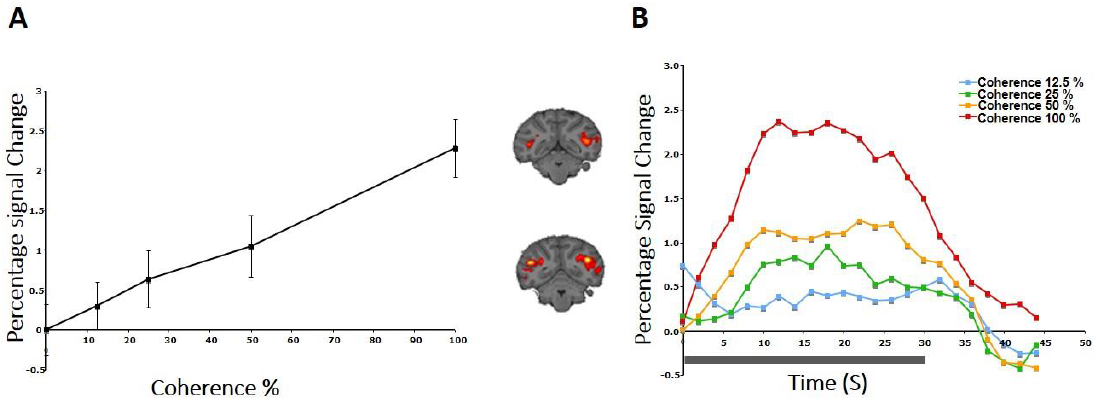
CBV response to coherence in area V5/MT+ of a rhesus macaque. *<u>A:</u>* Both area V5/MT and MST behaved in a similar manner and are grouped together. Note the monotonic increase of the CBV signal as a function of coherence. *<u>B:</u>* BOLD signal intensity modulation induced by presenting an RDK with motion coherence levels: 12.5%, 25%, 50%, 100%. Transition from the baseline (0% coherence) occurred at time 0 and the duration of the coherent RDK presentation was 30 seconds.

BOLD signal specific to motion coherence was present also in higher areas, as determined by contrasting all epochs of motion coherence against the 0% coherence baseline. Significant activation was observed in the cuneus (Cun) in 4/4 subjects (p<0.05, 1-way ANOVA), the superior temporal sulcus (STS) in 3/4 subjects (p<0.05, 1-way ANOVA), the posterior lateral sulcus (pLS) in 3/4 subjects (p<0.05, 1-way ANOVA) and the intraparietal sulcus (IPS) in 2/4 subjects (p=0.001, 1-way ANOVA). In STS, pLS and IPS, the mean BOLD magnitude increased as a function of motion coherence, similar to hV5/MT+. Each ANOVA measurement reflects mean response amplitude across trials at the same coherence level (fig. 5). Visual motion-related activation in these areas has been observed in earlier studies (Sunaert et al. 1999), except perhaps for pLS. We also observed significant modulation with coherence in the Cingulate (Cing) and Precuneus (PreC), but this was only for a single subject (1/4).

**Figure 5:**
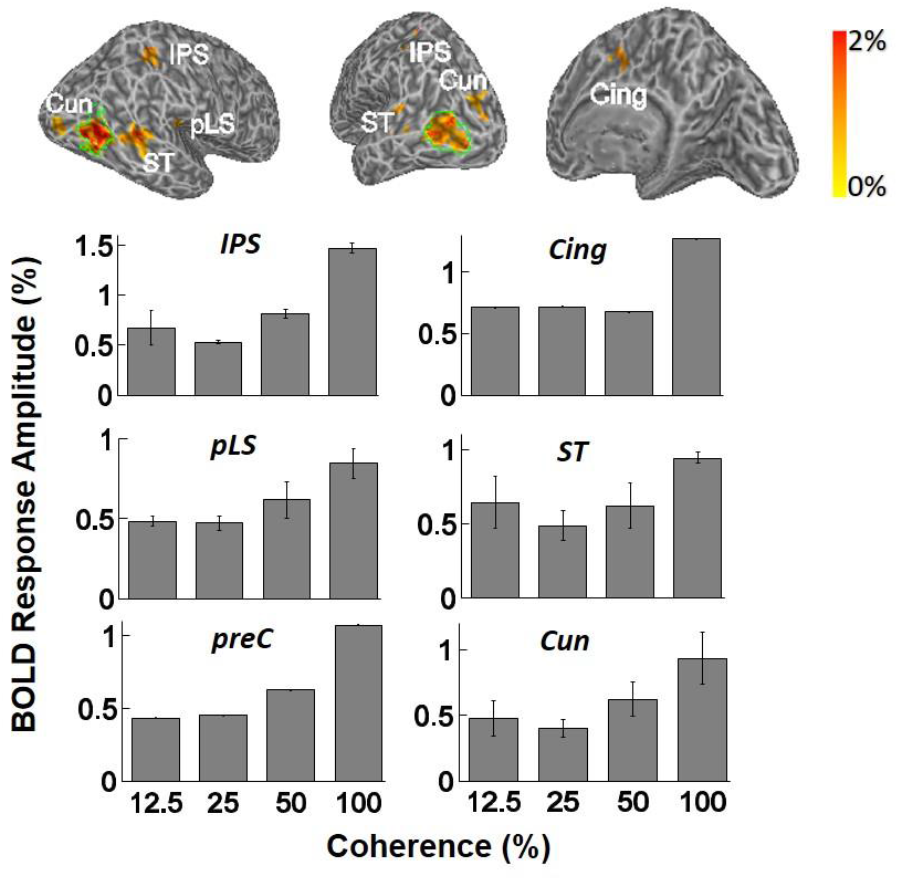
*Top:* examples of inflated white matter surfaces of individual subjects. Overlaid color maps represent the mean response amplitude across coherence levels of 25%, 50% and 100% and show only the region with corrected p values less than 0.05 in the GLM analysis. The green contours delineate the boundaries of area hV5/MT+ as determined from the hV5/MT+ localizer scans. Labels denote the anatomical locations of activated areas: ST-superior temporal sulcus, pLS – posterior lateral sulcus, Cun -cuneus, IPS -intraparietal sulcus, Cing -cingulate cortex, PreC – precentral sulcus. *Bottom: A*veraged fMRI response amplitudes in the activated brain areas (ST, IPS, Cing, PreC, pLS and Cun) across subjects. The mean BOLD magnitude increases significantly as a function of motion coherence. Only subjects that showed significant activation in these areas when we contrasted all-coherent RDKs against the baseline are averaged (please see result section for a discussion).

Results presented so far were obtained with passive stimulus presentation and did not involve performing a direction of motion discrimination task. In what follows, we compare visual responses while subjects are performing an RDK-dependent direction of motion discrimination task versus a luminance modulation task at fixation.

#### Performing direction of motion discrimination flattens motion-coherence dependence in hV5/MT+

##### Study B (bilateral stimulation)

Six additional subjects were tested with RDKs that were simultaneously presented in symmetric locations, one in each hemi-field. Subjects were asked to fixate versus to perform a direction of motion discrimination task in one of the hemifields. RDKs for this task were generated using the range-of-directions method introduced by Huxlin and Pasternak (Krystel R. Huxlin and Pasternak 2004), as we wanted to test coherence responses with the same stimuli used in visual rehabilitation (K. R. Huxlin et al. 2009). Four different coherence levels (range of motion: 0^0^, 180^0^, 288^0^, 360^0^) were tested, presented from a baseline range-of-motion condition of 360^0^ (no global motion coherence). As expected, both the left and the right hV5/MT+ complex exhibit increased modulation during epochs of stimulus presentation (fig. 6A-B). This was true even when the stimulus presented was the 360° RDK, which is identical to the baseline. This indicates that in this case the BOLD response does not reflect the characteristics of the stimulus itself, but rather stimulus anticipation and/or task-related effort (subjects were cued to respond to a new stimulus presentation by a change of color in the fixation spot; see methods). Interestingly, the BOLD signal response as a function of coherence differed between the “task relevant” versus the “task irrelevant” hemisphere. In particular, hV5/MT+ BOLD response was flat as a function of coherence in the task relevant hemisphere (the hemisphere contralateral to the RDK whose direction of motion the subject was tasked with reporting). In contrast, hV5/MT+ BOLD response increased as a function of coherence in the task irrelevant hemisphere, as observed in studies A and C (see below).

**Figure 6:**
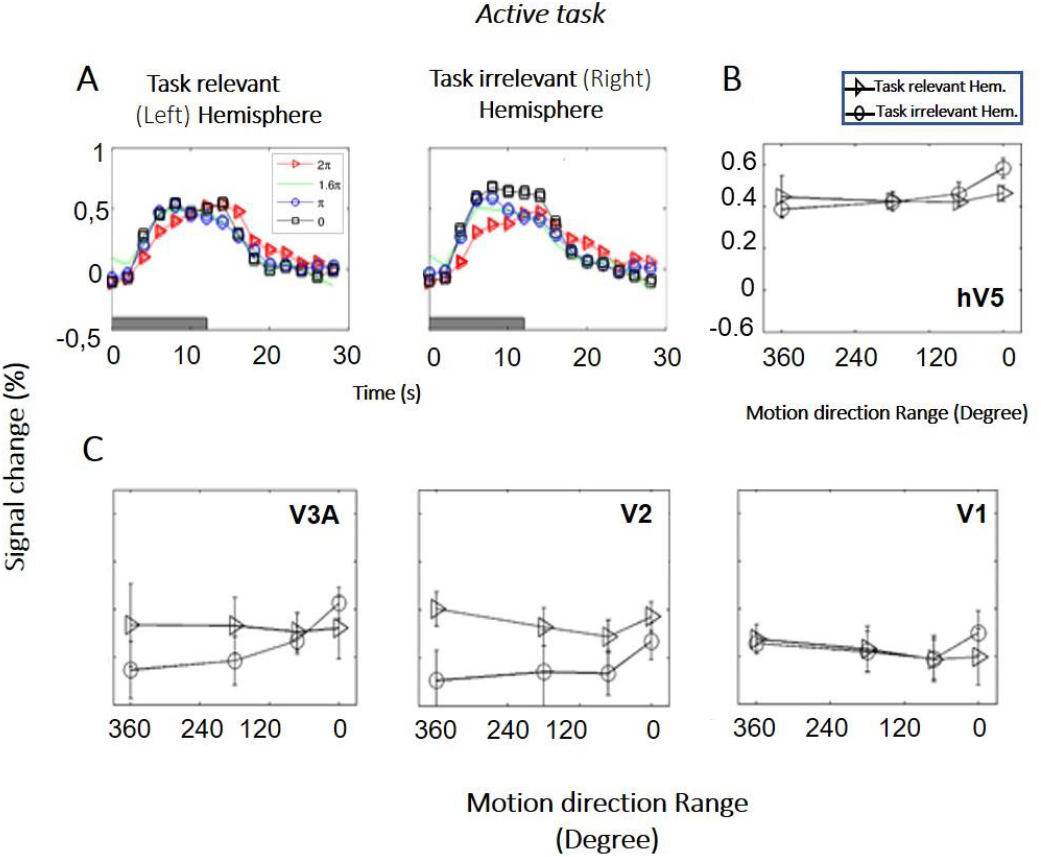
BOLD activity as a function of coherence in the task relevant vs the task irrelevant hemisphere (study B). **A**. BOLD signal change in hV5/MT+ of the task relevant (left) versus the task irrelevant (right) hemisphere of one subject. **B**. Average BOLD signal change in area hV5/MT+ of the task relevant (left, triangle) versus the task irrelevant (right, circle) hemisphere across 6 subjects. **C**. Same as B for areas V3A, V2 and V1. Error bars indicate standard error of the mean across subjects.

##### Study C (unilateral stimulation)

We used random dot stimuli to characterize responses in hV5/MT+ complex to motion coherence in six additional healthy subjects and seven patients under two conditions. In the first condition (passive task), subjects performed a fixation task at the center of the screen while RDKs of different coherences were presented unilaterally away from fixation (see Methods). In the second condition (active task), subjects were instructed to report the direction of motion of RDKs presented at identical locations, while maintaining fixation. RDKs for this task were generated using the range-of-directions method introduced by Huxlin and Pasternak (Krystel R. Huxlin and Pasternak 2004), as in study B. Note that RDK’s were smaller and unilaterally presented compared to RDKs in study A (which were 12 degrees in radius and centrally presented).

In the control subjects, the hV5/MT+ complex showed significant activation to motion coherence both in the contralateral hemisphere and in the ipsilateral hemisphere relative to the stimulus presentation. As expected, more voxels were significantly activated in the contralateral hV5/MT+ complex (fig. 7A), while voxels activated in the ipsilateral hV5/MT+ complex in part correspond to area MST, whose receptive fields are bilateral. Activated voxels showed strong BOLD signal modulation with coherence both ipsilaterally (presumably corresponding largely to area MST) and contralaterally with respect to the stimulus presentation (fig. 7B). Similarly to study A, hV5/MT+ increased with motion-coherence when the subject was performing a motion-unrelated task at fixation (fig. 7B; orange lines). However, when subjects were asked to perform direction of motion discrimination, the dependence of the BOLD signal on coherence was markedly suppressed and essentially completely abolished (fig. 7B; blue lines).

**Figure 7:**
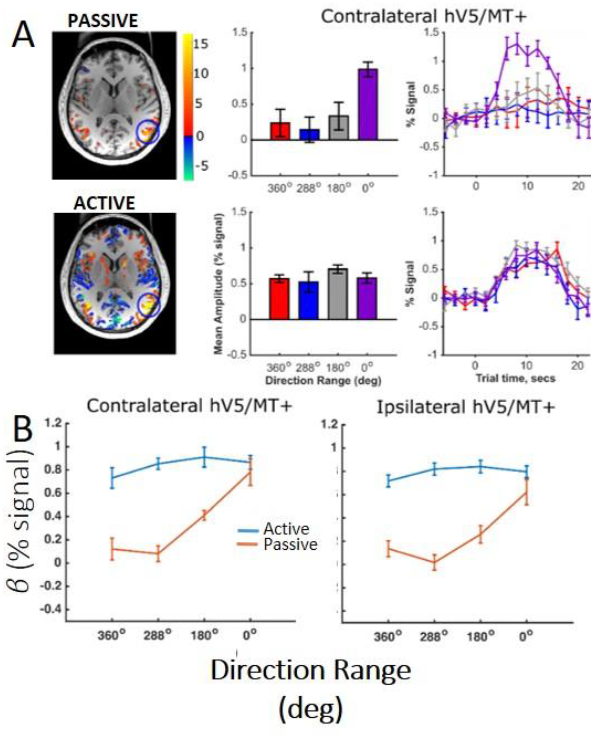
hV5/MT+ responses to motion coherence in control subjects (study C). ***A*:** Activation maps and average BOLD signal change in the contralateral hV5/MT+ of one subject, while the subject was performing an active direction of motion discrimination task (bottom) versus an unrelated task (passive) at fixation (top). For the passive condition our results are consistent with our previous findings and the literature that BOLD signal modulation increases with coherence when no discrimination task is employed. However, when the subject is performing direction of motion discrimination the BOLD signal modulation is approximately the same at all coherences. ***B*:** Average % BOLD signal change for all participants (N=6) performing the motion discrimination task (blue line) versus fixation (orange line). Error-bars indicate standard error of the mean across subjects. As in (A), when the subjects are performing the direction of motion discrimination task the BOLD signal modulation is approximately the same at all coherences. In contrast, when subjects are fixating without performing the motion discrimination task BOLD signal modulation increases monotonically with coherence level. A similar activation pattern is also observed in ipsilateral hV5/MT+, presumably reflecting voxels that belong primarily to MST whose receptive fields are bilateral.

### Patients with dense cortical visual field scotomas

We measured the hV5/MT+ responses to motion coherence for 7 subjects with early visual cortical lesions resulting in a dense homonymous visual field scotoma (see methods). Three patients (patients S04, S07 and V1003) were tested while performing the *passive fixation task*. Six patients, two of which had performed the passive task (S04, V1003) and four other patients (S12, S15, S29, S02), were tested while performing the *active task*. Patient responses were tested both in the sighted and in the blind visual field.

#### Responses in the sighted visual field

*P*atients did show significant activation in both the contra-lesional and ipsi-lesional hV5/MT+ when the stimulus was presented in their *sighted visual field* (fig. 8A.ii; fig. 9B). The dependence of BOLD responses in both contra- and ipsi-lesional hV5/MT+ to motion coherence was similar to the healthy subjects presented above, i.e. linearly increasing with coherence when subjects were performing a task at fixation (fig. 8B) and flat when subjects were engaged in a direction of motion discrimination task (fig. 9C). As expected, the patients’ performance in the direction of motion discrimination task was commensurate to control subjects when the stimulus was presented in the sighted field (fig. 9A).

**Figure 8:**
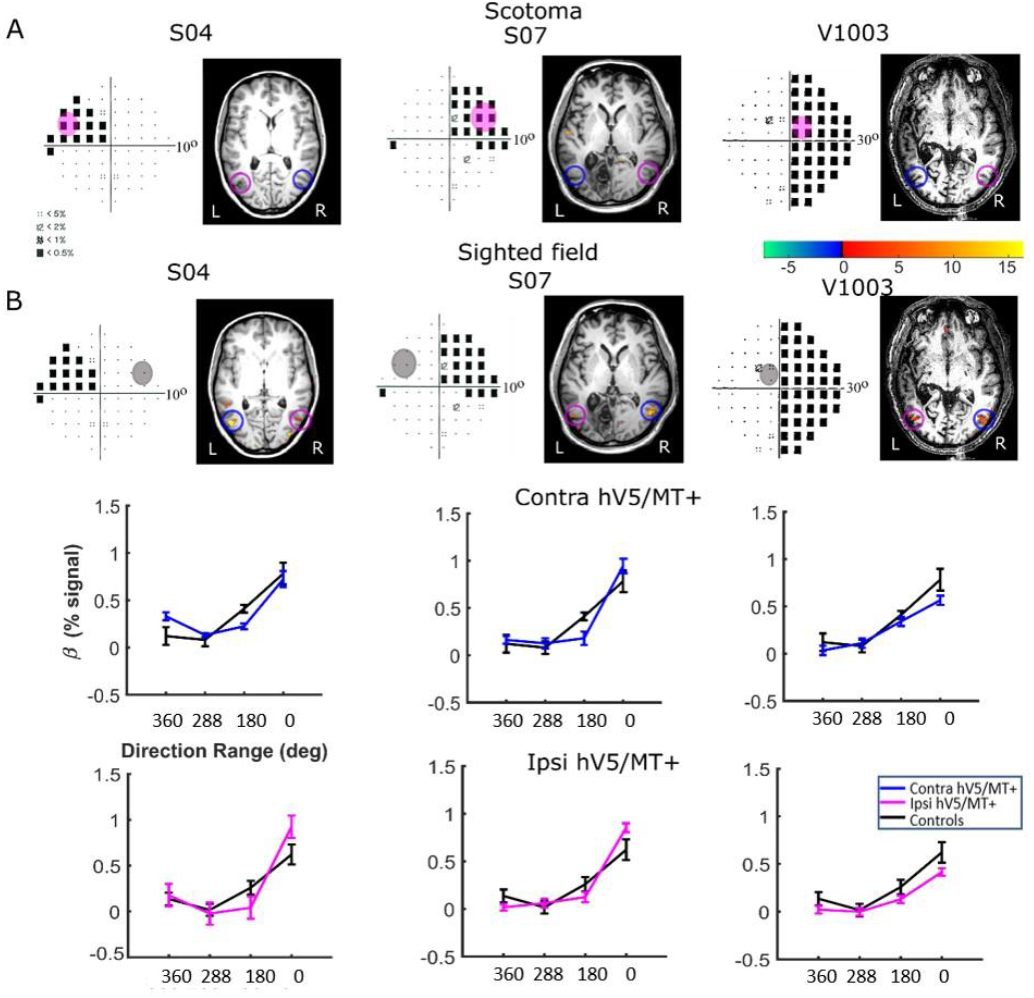
hV5/MT+ responses to motion coherence in three patients during the passive task. ***Ai:*** Visual field deficits for each patient based on perimetry and activation maps when the stimulus was presented within the blind visual field (magenta disk). We found no significant activity in either the contralesional (blue circle) or ipsilesional (magenta circle) hV5/MT+ when the stimulus was presented within the patients’ scotoma and the patients were performing a passive fixation task. ***A*.*ii:*** Activation maps when the stimulus was presented within the sighted visual field (grey disk). We found significant activity for both the contralesional (blue circle) and ipsilesional (magenta circle) hV5/MT+ when the stimulus was presented in the sighted field of the patients. ***B:*** mean GLM *β* weights of hV5/MT+ across patients as a function of coherence level when the stimulus was presented within their sighted visual field compared with control subjects (black). Responses in both the contralesional (blue) and ipsilesional (magenta) hV5/MT+ were similar to control subjects.

**Figure 9:**
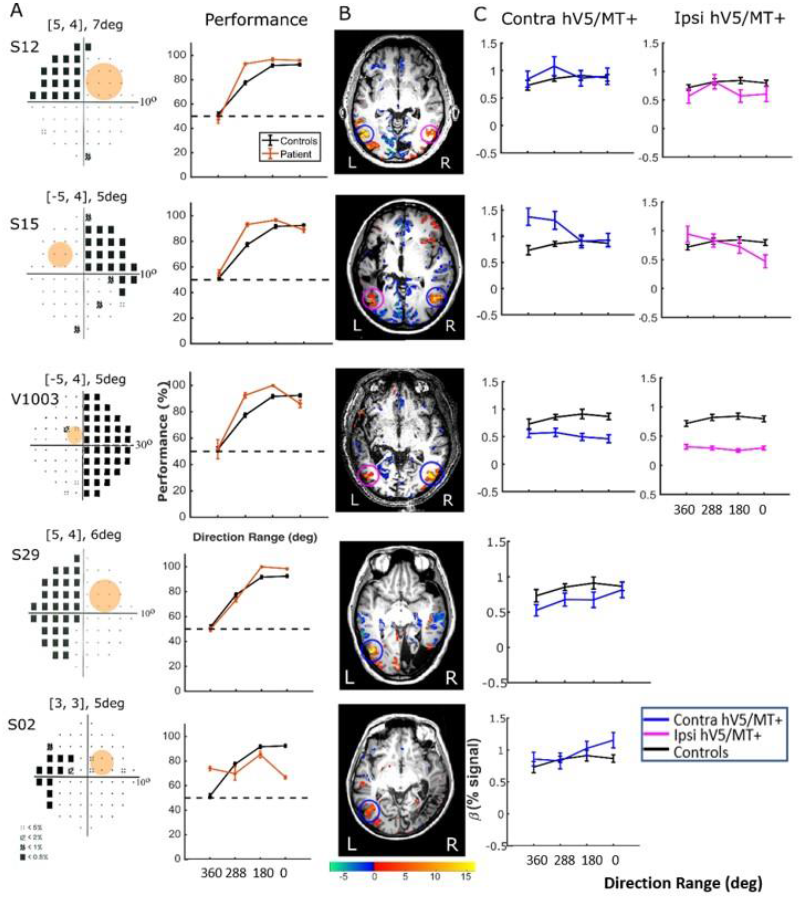
Behavioral performance and hV5/MT+ responses to motion coherence within the sighted field of patients performing a motion direction discrimination task. ***A:*** Left column: Visual field deficits for each patient based on perimetry and location of the stimulus aperture (orange disk). Right column: behavioral performance for the motion direction discrimination task when the stimulus is presented within the sighted field of patients (orange) versus control subjects (black). ***B:*** Activation maps when the stimulus was presented within the sighted visual field. We found significant activity for both the contralesional (blue circle) and ipsilesional (magenta circle) hV5/MT+. ***C:*** mean GLM *β* weights of contralesional (blue) and ipsilesional (magenta) hV5/MT+ across patients as a function of coherence level compared to control subjects (black).

#### Responses in the blind visual field

Although retinotopic mapping did reveal visually driven activation in both ipsi-lesional and contra-lesional hV5/MT+ contrasting coherently moving stimuli to stimuli of 360°- direction of motion range, i.e. no motion coherence, revealed no significant activation during the passive fixation condition when the RDK stimulus was presented inside the patient’s scotoma (fig. 8A.i). Note that this does not necessarily mean that hV5/MT+ of these patients does not get activated by motion stimuli presented within the scotoma, but rather that the modulation is flat as a function of coherence (i.e. it does not differ significantly from the 360°-range coherence baseline). Therefore, we can conclude that hV5/MT+ RDK responses arising from stimuli presented inside the dense cortical scotoma, are either too weak to elicit significant modulation and/or do not vary significantly as a function of motion coherence.

In contrast, performing the direction-of-motion discrimination within the patients’ scotoma elicited significant responses in hV5/MT+, as judged by contrasting all RDK coherent-motion conditions tested against the baseline (360°-direction of motion, i.e. no motion coherence) (fig. 10).The BOLD signal as a function of coherence was suppressed and approximately unchanged at all coherences similar to the responses observed in healthy subjects and patients when the stimulus was presented in their sighted field under the direction of motion discrimination condition. This is markedly different than the profile of BOLD signal modulation as a function of coherence during the passive fixation condition when the stimulus was presented in the sighted field (fig. 8B). This occurred while the patients’ performance remained at chance at all coherence levels when the stimulus was presented in their blind field, confirming the visual field defect, and also suggesting that the subjects did not significantly break fixation (fig 10A).

**Figure 10:**
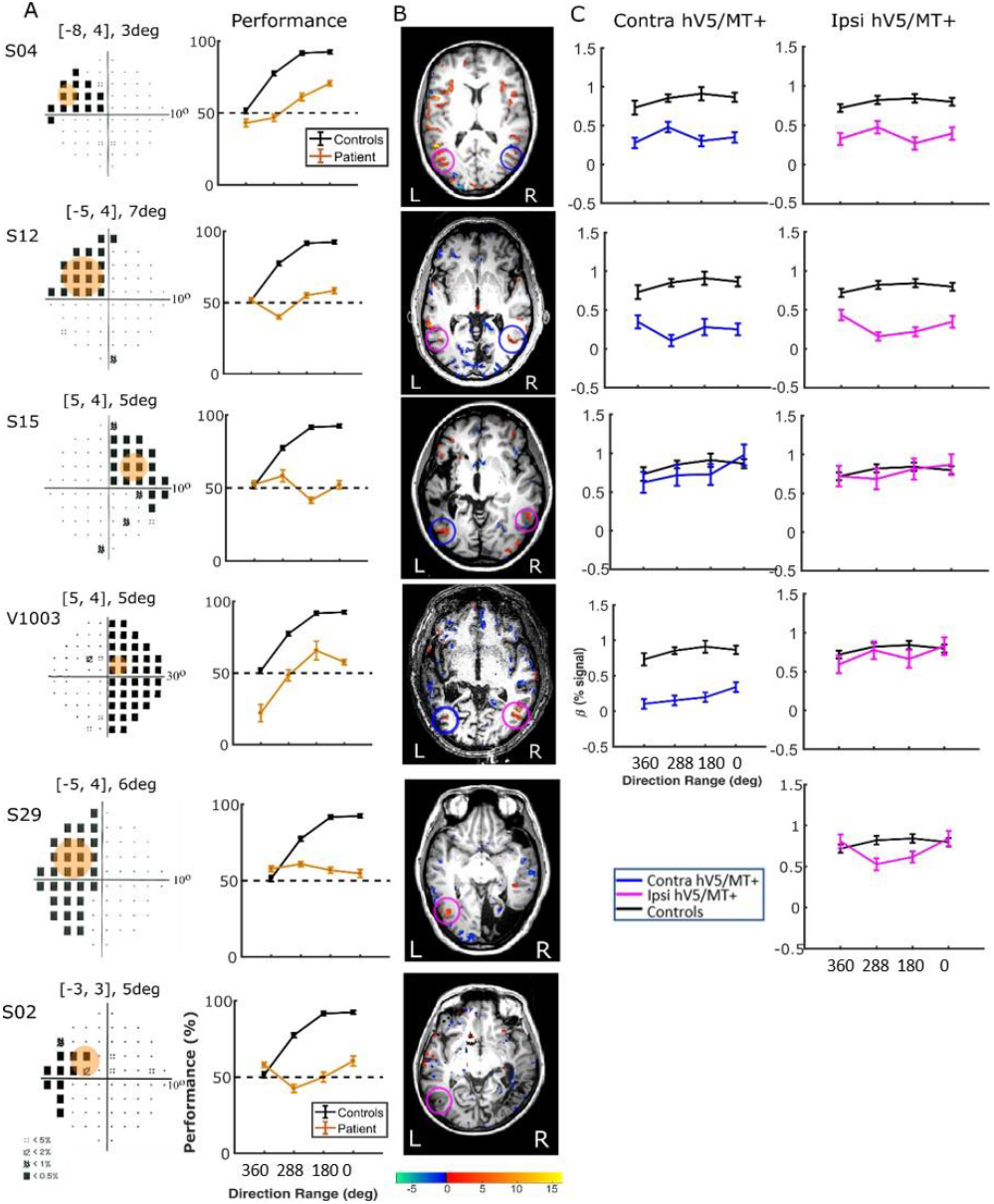
Behavioral performance and hV5/MT+ responses to motion coherence within the blind field of patients performing a motion direction discrimination task. ***A:*** Left column: Visual field deficits for each patient based on perimetry and location of the stimulus aperture (orange disk). The center coordinates and diameter of the aperture are shown on top of each graph in the form [x, y], zdeg, where x=deg from vertical meridian, y=deg from horizontal meridian and z=deg diameter. Right column: behavioral performance for the motion direction discrimination task when the stimulus was presented within the blind field of patients (orange) versus control subjects (black). ***B:*** Activation maps when the stimulus was presented within the blind visual field. We found significant activity for both the contralesional (blue circle) and ipsilesional (magenta circle) hV5/MT+. ***C:*** mean GLM *β* weights of contralateral (blue) and ipsilateral (magenta) hV5/MT+ across patients as a function of coherence level compared to control subjects (black).

There were some differences across subjects, but these do not change the basic observations reported above. Specifically, for patients S02 and S29, hV5/MT+ was itself lesioned and only ipsi-lateral data could be obtained (all other patients showed bilateral hV5/MT+ activation upon unilateral stimulus presentation). The hV5/MT+ area activated contralateral to the stimulus presentation was significantly smaller than control subjects

## 6 Discussion

This study revisited the question of how human visual cortex responds to motion coherence stimuli. We went further than previous studies in several respects: 1) we demonstrated how BOLD fMRI responses are modulated by task relevance, 2) we used the non-coherent motion condition as baseline instead of a static stimulus in order to maximize specificity to global motion integration, 3) we validated part of our findings in a non-human primate, and 4) we showed that our observations remain consistent in patients with visual scotomas arising due to cortical injury.

Overall, in healthy subjects, passive RDK presentation starting from a baseline condition of 0% coherence resulted in robust activation of the hV5/MT+ complex and area V3A, as well as the higher areas (Cun, pLS, ST, Cing,IPS,preC). The result in hV5/MT+ complex is in general agreement with (Rees, Friston, and Koch 2000; Becker, Erb, and Haarmeier 2008) whose baseline condition involved static stimuli. On the face of it, the amplitude of the BOLD response to 100% coherence in area hV5/MT+ we observe (∼0.5%) appears to be stronger than the one reported by Rees et al. (0.1%), but this difference can be largely explained on the basis of a difference in the experimental design. Specifically, Rees et al used an event related paradigm with much shorter stimulus presentation periods. Once this difference is taken into account, one arrives at very similar underlying firing rate estimates using the model introduced by Rees et al. (Rees, Friston, and Koch 2000). This result is reassuring, as the baseline condition we used, unlike the one selected by Rees et al. and Becker et al., does maximize the chance that the observed BOLD responses are selective for the integration of local motion signals to global motion coherence. It is also consistent with Braddick et al. (Oliver J. Braddick et al. 2001) who had also used a dynamic moving baseline with 0% coherence obtaining similar results.

On the other hand, our results are in contrast with results by Smith et al., and McKeefrey et al. (Smith et al. 2006; McKeefry et al. 1997) who found that area hV5/MT+ responses were not modulated by motion coherence. We believe that differences in the experimental conditions are the most likely reason for the disagreement in conclusions. Specifically, Smith et al. and McKeefrey et al. employed paradigms with very low RDK dot densities leading to a decreased chance for motion integration within the receptive fields of neurons in hV5/MT+. The BOLD response in each voxel depends not only on the strength of the motion signal but also on the number of different motion directions stimulated within its population receptive field. When motion coherence increases, the number of different motion directions stimulated decreases. As first pointed out by Braddick et al. (Oliver J. Braddick et al. 2001), the balance between these two factors changes with dot density. For example, accounting for the eccentricity of our stimuli we calculated that the average number of moving dots within the typical area of an hV5/MT receptive field would be about 50, compared with about 4-6 for the stimuli used by (Smith et al. 2006; McKeefry et al. 1997). Therefore, the number of dots in these two studies is much lower than our study and also than Braddick et al. (Oliver J. Braddick et al. 2001), Rees et al. (Rees, Friston, and Koch 2000) and Becker et al. (Becker, Erb, and Haarmeier 2008). Hence, when the level of coherence increases, the increased global motion strength may not be able to overcome the decreased number of motion directions that stimulate the typical area hV5/MT voxel. The degree of motion opponency in different areas likely also plays a role.

Another noteworthy finding of our study is that unilateral RDK stimulus presentation modulated both the ipsilateral and contralateral hV5/MT+ (fig. 7B). Although the number of ipsilateral voxels activated in the hV5/MT+ complex were fewer than the contralateral ones, presumably corresponding to area MST whose large receptive fields cross the midline, we found that their coherence dependence was similar to those in the contralateral side. This suggests that coherence dependence is similar for both areas V5/MT as well as MST. Moreover, in a separate analysis (fig. 3C) separating area hV5/MT+ voxels to anterior and posterior groups did not reveal a difference in coherence dependence (fig. 3C), supporting that there is no strong difference in coherence dependence between area hV5/MT and the more posteriorly located, putative MST. This agrees qualitatively with Becker et al. (Becker, Erb, and Haarmeier 2008), who found area MST responses increase with motion coherence.

A novel and surprising finding in our study was the suppression of coherence dependence in areas hV5/MT+ and V3A observed when healthy subjects performed a direction of motion discrimination task instead of passive fixation. We conjecture that increased effort and/or allocated attention during the active visual motion discrimination task, likely increases disproportionately the BOLD responses in these motion selective areas for low coherences, thus flattening the coherence dependence (see figures 6B and 7B). BOLD signal responses in our study were increased commensurately even for the non-coherent stimulus condition, which was identical to the baseline, suggesting that they are not specific to global motion integration across space. It is thus probable that coherence dependence is masked by a non-selective effort-related increase in the BOLD signal response within fMRI voxels. Attention to the stimulus or effort during the motion discrimination task would tend to increase the BOLD response in hV5/MT+ (Huk, Ress, and Heeger 2001) and this may indeed be the underlying reason for our findings. It remains to be investigated whether ensuring that the load of attention or effort remains the same across different levels of coherence, might reinstate the monotonic dependence of hV5/MT+ activation as a function of coherence.

### Motion coherence responses in areas outside the hV5/MT+ complex

We also found modulation of the BOLD signal as a function of coherence in higher motion-processing areas, such as the cuneus, superior temporal sulcus, posterior lateral sulcus, and the intraparietal cortex. Others have also documented parietal activation in response to discrimination of motion (Orban, Van Essen, and Vanduffel 2004; Peuskens et al. 2004; Vanduffel et al. 2002) as well as in area hV5/MT+ and lateral occipital cortex (O. J. Braddick et al. 2000; Grill-Spector, Kourtzi, and Kanwisher 2001; Kourtzi and Kanwisher 2001). BOLD signal in these areas increased with motion coherence qualitatively similarly to area hV5/MT+ (fig. 3).

In contrast, responses in early visual areas (V1/V2) decrease as coherent motion strength increases (fig. 3B). This finding agrees with earlier reports by Braddick et al. (Oliver J. Braddick et al. 2001), but contrasts with McKeefry et al. (McKeefry et al. 1997) who found the opposite. Again, this is likely the result of the difference in the density of the dots between our stimuli. At 6° eccentricity, the mean receptive field size of area V1 neurons is 0.53 (Dow et al. 1981), so the average number of dots within a typical V1 receptive field would be ∼0.5. Therefore, increased coherence level does not enhance the motion signal strength detected by individual V1 neurons. The BOLD response in V1 would then be solely determined by the number of different motion directions that fall over time in the receptive field. As coherence level increases from 0% to 50%, the density of dots corresponding to random noise changes from 2/degree^2^ to 1/degree^2^, and the number of motion directions stimulating each voxel over time varies relatively little. However, at 100% coherence, the number of motion directions corresponding to random noise is reduced fairly abruptly to zero. This may explain the relative flat dependence of V1 BOLD responses at coherence levels of 12.5%, 25%, and 50%, versus the prominent negative response at coherence 100% (fig 3A; passive task). Interestingly, when subjects perform a direction of motion discrimination task, the response to coherence is flat or weakly positive at all coherence levels (fig. 6C). As discussed above, this is likely a result of an increase in effort and/or attention to direction of motion signals.

Consistent with our findings in healthy controls, when the stimulus was presented passively in the non-lesioned hemisphere of patients with V1 injury (fig. 8A.ii), we found that both contra- and ipsi-lesional hV5/MT+ were activated and showed monotonically increasing motion coherence dependence similar to the controls (fig. 8B). Bilateral hV5/MT+ activation with unilateral stimulus presentation in the seeing field, has also been reported in other studies, such as in Ajina et *al*. (Ajina et al. 2015), who also showed that the presence of the contra-lesioned (intact) V1 is important for promoting the communication between the two hemispheres. Performing the direction-of-motion discrimination task when the stimulus was presented in the patients’ sighted field (fig. 9A) again elicited responses commensurate to healthy subjects, suppressing coherence dependence. When the stimulus was presented passively in the blind hemifield of our patients hV5/MT+ activation did not reach significance, in agreement with Ajina et al. (Ajina et al. 2015). However, performing the direction-of-motion discrimination task within the scotoma was sufficient to induce significant hV5/MT+ activation (fig.10), even though subject performance remained at chance. Again, the strength of induced activity did not depend on motion coherence and is likely effort related (Masuda et al. 2021).

### Eye movements do not explain the effects we observed

Although subjects were trained to fixate at the fixation mark at the center of the stimulus, the possible presence of pursuit eye movements and variable attentional shifts cannot be completely excluded. However, it is very unlikely that our results can be explained on the basis of aberrant eye-movements. First, during the fixation-only task subjects maintained high performance (> 96%) on a rigorous task at fixation, suggesting they fixated well. Second, the effect of pursuit eye movements is expected to be higher at high motion coherence, yet, across subjects, responses at fully coherent RDKs were similar between the fixation-only condition (during which subjects fixated well) versus the motion discrimination condition. Furthermore, in study C, experiments were performed under eye-tracking and post-hoc analysis confirmed that subjects maintained fixation. Presenting the stimuli in the sighted field of our patients shows similar results with the healthy subjects, increasing confidence that the patients were also fixating well and the basic observations seen in healthy controls are also true for the healthy cortex of the patients.

In summary, in agreement with the literature, we found that BOLD responses in area hV5/MT+ were monotonically increasing when subjects did not actively perform a motion discrimination task. In contrast, when subjects performed an RDK direction of motion discrimination task, hV5/MT+ BOLD responses became flat as a function of coherence, probably as a result of increased attention or task-related effort load at low coherences. The same effect was observed when RDK stimuli were presented in the sighted field of the patients. When the stimulus was presented inside the patients’ scotoma, performing the motion discrimination task was necessary in order to observe significant hV5/MT+ activation. However, hV5/MT+ activation was again not stimulus coherence dependent, likely representing top-down mediated task dependent effects as argued by (Masuda et al. 2021). In general, these observations shed further light on how visual cortex responses behave as a function of motion coherence, preparing the ground for using these methods to study visual system recovery after injury.

## 9 Author Contributions

AR integrated the data and together with AP wrote the final form of the manuscript. XZ and DP wrote a first draft of the manuscript. XZ, AP, GK performed the experiments. AR, AP, GK analyzed the fMRI data. SMS, AR and GK contributed to conception and design of the study. AP was only involved in Study C. All authors contributed to the manuscript revision, read, and approved the submitted version.

## 11 Abbreviations

hV5: human Visual 5 area
MT+: middle temporal area complex
RDKs: Random dot kinematograms
fMRI: Functional magnetic resonance imaging
BOLD: Blood Oxygen Level Dependent

## 12 Acknowledgments

We thank Xinmiao Peng and John Arsenault for their help with experimental work in Study A and in monkey study, respectively.

## 13 Contribution to the field statement

Random dot kinematogram (RDK) stimuli have been recently introduced for rehabilitating visual motion perception of patients with lesions of the visual cortex. It is important to understand the responses of visual areas to such stimuli in order to use them for studying the recovery of visual perception. Here, we measured visual area responses to such stimuli in normal subjects. In agreement with prior studies, we found that visual area responses change as a function of RDK motion strength (motion coherence), when subjects are fixating. However, this dependence is abolished when subjects are engaged in performing a direction of motion discrimination task, likely as a result of differential attention or task-related effort load. Presenting the stimuli in the sighted field of patients showed similar results with the healthy subjects increasing confidence that the patients were also fixating well and that the basic observations seen in healthy controls are also true for the healthy cortex of the patients. We discuss possible reasons for our findings in the context of prior literature. These observations shed further light on how normal visual cortex responses behave as a function of motion coherence, preparing the ground for using these methods for studying visual system recovery after injury.

## 15 Data Availability Statement

Datasets are available on request: The raw data supporting the conclusions of this article will be made available by the authors, without undue reservation.

## Footnotes

1 This coherence is not related to the motion stimulus

